# Robotic Handling Preserves Induced Pluripotent Stem Cell Derived Vascular Smooth Muscle Cells Differentiation using a Weekend-free, Automatable, Low-variability, Low-cost (WALL) Protocol

**DOI:** 10.1101/2025.11.25.690512

**Authors:** Monique Bax, Jeya Ramalingam, Valentin Romanov, Muhammad A Alsherbiny, Keerat Junday, Jordan Thorpe, Emily Hurley, Melanie Lim, Ingrid Tarr, Stephanie Hesselson, Eleni Giannoulatou, Renjing Liu, Adam Hill, Siiri Iismaa, Robert M Graham

**Affiliations:** Victor Chang Cardiac Research Institute, Darlinghurst, NSW, 2010; St Vincent’s Hospital, Darlinghurst, NSW, 2010; School of Clinical Medicine, St Vincent’s Healthcare Clinical Campus, Faculty of Medicine and Health, UNSW Sydney, Australia

**Keywords:** Induced Pluripotent Stem Cells, Vascular Smooth Muscle Cells, Tissue Engineering, Proteomics, Liquid Handling Robotics, High-throughput, Drug Screening

## Abstract

Vascular disease modeling and tissue engineering require scalable, efficient, and reproducible methods for differentiating cells, however no studies have validated whether automation, essential for scale-up, preserves differentiation quality in cells. Here, we developed a differentiation protocol for induced pluripotent stem cell (iPSC)-derived vascular smooth muscle cells (iVSMCs) that is applicable to both manual, and automated robotic culture. iPSC lines generated from male and female donors across various ages were used to optimize the generation of iVSMCs. iVSMCs had consistent morphology and protein expression was largely similar to primary VSMCs, including expression of myosin heavy chain-11 (MYH11), α-smooth muscle actin (α-SMA), transgelin (TAGLN) and calponin (CNN1). Functionally, these iVSMCs could respond to the vasoconstrictor carbachol. Coupling this protocol with an automated Hamilton liquid-handling robotics system allowed the generation of iVSMCs in large quantities, morphologically similar to manually differentiated iVSMCs. Comparative proteomic analysis confirmed protein expression did not differ between automated and manual differentiation methods. Automation markedly reduced manual labor and facilitated increased production without sacrificing cell quality. This study demonstrates the feasibility of automating iVSMC differentiation, marking a significant step towards scalable VSMC manufacturing for three-dimensional applications, and for the ever-growing demands of organoid and tissue engineering applications.

**Statement of significance:** The rapidly expanding fields of tissue engineering and high-throughput disease modeling require large quantities of viable cells, making automation essential. Robotic liquid handling systems are increasingly employed for generating large quantities of induced pluripotent stem cell (iPSC)-derived cells, requiring the development of scalable protocols. No studies have yet evaluated whether the use of robotic liquid handling alters the differentiation of cells compared to traditional manual handling. Here, we report a scalable differentiation protocol that robustly generates iPSC-derived VSMCs (iVSMCs) from a cohort of female and male donors of varying age (n=5 lines). We demonstrate for the first time that iVSMCs, differentiated through an automated robotic pipeline, had no significant differences in morphological or proteomic phenotype compared to manual differentiation. The ability to automate this protocol facilitates robust and reproducible differentiation of large quantities of iVSMC, setting a precedent for the application of automation in generating iPSC-derived cells.

**Graphical Abstract:** 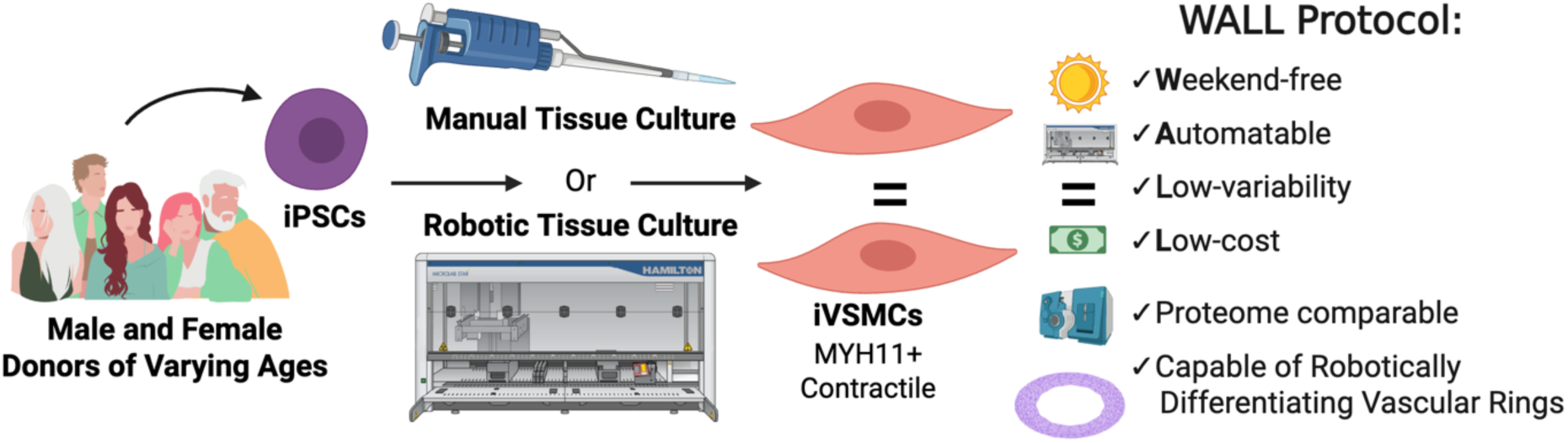

## Introduction

Cardiovascular diseases are projected to cost $1.1 trillion by 2035 in the US alone, with blood vessel-associated conditions including high blood pressure and coronary heart disease accounting for, or contributing to, the overwhelming majority of cases^[1]^. Vascular smooth muscle cells (VSMCs) are a central component of blood vessels, responsible for wall contractility, regulating blood flow and blood pressure ^[2]^, and are accordingly a central focus of efforts to improve vascular health. As the prevalence of these vascular-linked diseases continues to increase globally, so too does the demand for advanced VSMC research, such as drug screens and bioengineered products for regenerative medicine applications^[3]^, which require large volumes of VSMCs.

VSMCs present unique challenges for *in vitro* modeling. Unlike cardiac muscle cells (cardiomyocytes) or skeletal muscle cells, VSMCs are not terminally differentiated^[4]^. This plasticity allows them to oscillate between contractile and synthetic states to facilitate normal vascular growth and repair, but also leaves these cells vulnerable to the pathogenesis of a myriad of vascular disorders, including atherosclerosis and arterial dissections^[5–7]^. Synthetic VSMCs are characterized by active cell proliferation, migration, and extracellular matrix (ECM) production^[8,9]^. Fluid dynamics is a key driver for phenotype switching in VSMCs, which sense altered pressure, and turbulence^[10,11]^. To best model this dynamic behavior both in the context of development, as well as in health and disease, induced pluripotent stem cell-derived VSMCs (iVSMCs) are increasingly employed, capturing the diverse genetic backgrounds of individuals in a human-derived system^[3,12]^.

Current induced pluripotent stem cell (iPSC) differentiation methods to generate VSMCs have provided invaluable insights into vascular biology and disease mechanisms, examining both the effects of the embryological backgrounds of the cells and phenotypic variations^[13]^. However, challenges remain for the field, including a lack of information regarding the effects of scalability and automation, a high cost of implementation, particularly labor costs, as well as reagent costs, and limited reporting on the robustness of protocols across multiple lines^[12]^.

Scalability requires a highly efficient differentiation protocol to generate a consistent cell type, with low batch-to-batch variability, and low phenotypic variability. Automating liquid handling represents an attractive option to address standardization in tissue culturing through minimization of human variation, and errors in handling; to reduce institutional liability through limiting exposure of staff to compounds that are potentially toxic; to reduce labor costs, and to maximize staff productivity^[15]^. However, to our knowledge, the impact of automated liquid handling on differentiation has not yet been examined for any cell type. For VSMCs, which alter their cellular identity in response to fluid dynamics^[16,17]^, determining if robotic liquid handling impacts differentiation and maintenance of iVSMCs is as an important step in assessing its appropriateness for producing cells in large scale via automation.

Recognizing these critical gaps, we report a novel differentiation protocol that addresses the urgent need for robust, functionally relevant iVSMCs. This protocol is designed to be easy to use, with low-variability and low-cost, as well as being weekend-free, which significantly enhances the reproducibility and accessibility of iVSMCs generation. Using multiple iPSC lines generated from male and female donors of varying ages, we show that iVSMCs can be generated both by manual handling (traditional tissue culture), as well as by automated means. The resultant cells expressed hallmark VSMC proteins, including myosin heavy chain-11 (MYH11), α-smooth muscle actin (α-SMA also known as ACTA2), transgelin (TAGLN also known as SM22) and calponin (CNN1). The iVSMCs produced also contracted in response to the cholinergic agonist, carbachol and, importantly, proteomic analysis of cells produced either manually or by the automated protocol showed no phenotypic differences in terms of the protein pathways expressed. As proof-of-principle for the use of the cells in future applications, we demonstrate that they can form three-dimensional vascular rings. The current protocol developed to automate iVSMC differentiation, paves the way for their scalable generation, serving to improve disease modeling, drug discovery, and tissue engineering to combat vascular disease.

## 1. Results and Discussion

### 1.1 Generation of Novel Healthy Female and Male Donor-Derived iPSC Lines to Validate Differentiation Protocol Robustness

To evaluate the robustness of our differentiation protocol, we generated a cohort of induced pluripotent stem cell (iPSC) lines from both female (n=3) and male (n=2) donors across a broad age range. One iPSC line (Line 4; SCVI15) was kindly provided by the Stanford Cardiovascular Institute Biobank^[18]^, while all other lines were successfully reprogrammed from fibroblasts or peripheral blood mononuclear cells (PBMCs) using Sendai virus as previously described^[19–21]^. All lines were subjected to a characterization pipeline (**Figure 1a**). The cell donors of this cohort included female and males ranging from 16-71 years of age at time of collection (**Figure 1b**). The pluripotent status of the reprogrammed lines was confirmed via immunostaining by the pluripotency markers OCT4 and SSEA4 (**Figure 1c**). The capability for trilineage differentiation was also demonstrated through immunocytochemistry; the potential of these iPSC lines to differentiate into cells expressing hallmark proteins, including PAX6, TAGLN and SOX17, which are indicative of successful differentiation to ectoderm, mesoderm, and endoderm, respectively (**Figures 1d-1f**). Gene expression profiles were analyzed using the hPSC Scorecard to further validate that these iPSC lines were capable of self-renewal and trilineage differentiation (**Figure 1g**).

**Figure 1.**
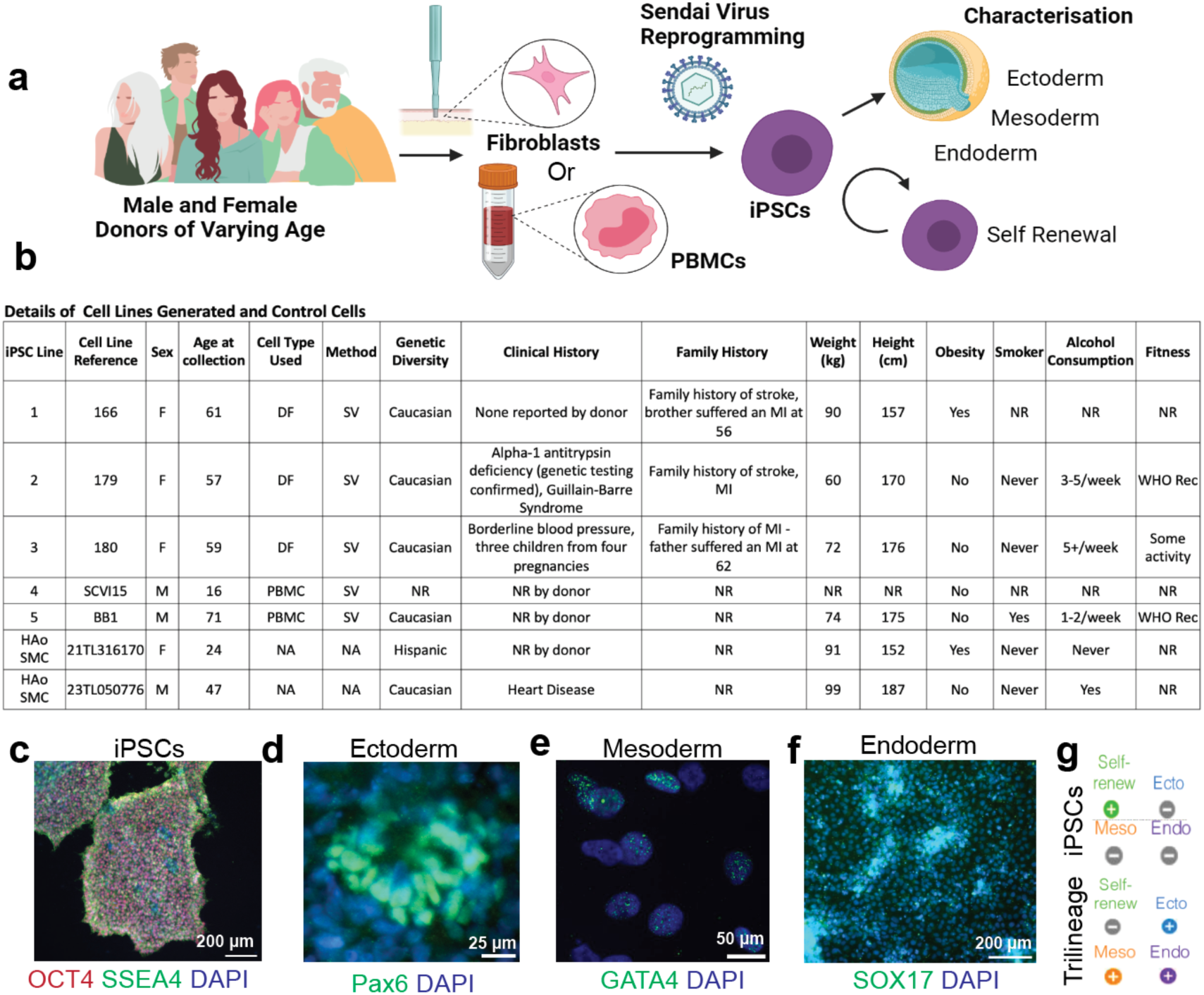
Reprogramming iPSC lines from healthy donors of diverse age and sex. **a)** Schematic and **b)** details of iPSC lines that were generated from fibroblasts or PBMCs from male and female donors of various ages and with differing health statuses using Sendai virus reprogramming. iPSC: induced pluripotent stem cell, HAoSMC: Human Aortic Smooth Muscle Cells, F: Female, M: Male, NR: Not Reported, MI: Myocardial Infarct, WHO Rec: World Health Organization recommended levels of exercise (**c**) Representative immunofluorescence images show pluripotency marker expression (OCT4, red; SSEA4, green; nuclear stain, blue). Immunocytochemistry following trilineage differentiation demonstrates capacity for three germ layer differentiation**: d)** PAX6 (green; DAPI nuclear stain, blue) expression indicates ectoderm differentiation, **e)** TAGLN (green; nuclear stain, blue) expression indicates mesoderm differentiation, and **f)** SOX17 expression indicates endoderm differentiation. **g)** Representative hPSC Scorecard analysis depicting the self-renewal capabilities and differentiation potential that were further validated by RNA microarray analysis of the lineage-specific differentiation potential of the iPSC lines.

To ensure our differentiation protocol was robust and broadly applicable, we prioritized diversity in donor age and sex when generating this cohort of iPSC lines. The inclusion of both female and male lines in studies is particularly important, as vascular cells are known to exhibit sex-specific differences from birth, regardless of hormones^[22,23]^. Moreover, because some epigenetic signatures of aging can be retained following reprogramming^[24]^. Together, diversity across these two parameters enabled the evaluation of differentiation efficiency in a biologically relevant cohort. All lines in this initial cohort were derived from individuals of Caucasian genetic diversity (except line SCVI15 which was deidentified), reflective of the European ancestry majority in Australia^[25]^. While this enabled certain baseline comparisons, we acknowledge the lack of ethnic diversity as a limitation. In future studies we aim to broaden ethnic representation to better reflect the global population and to ensure inclusivity.

### 1.2 Robust iVSMC Differentiation

Having validated our iPSC lines, we sought to develop a weekend-free, automatable protocol that was relatively Low cost, and low variability (WALL) that could generate differentiated cells at scale. This design philosophy addresses the practical barriers that limit VSMC production in both academic and industrial settings. A range of variables including media composition, dissociation methods, cell seeding density, and maturation time were trialed that ultimately resulted in a robust, monolayer-based protocol that can be generated under xenogenic-free conditions (**Figure 2a**).

**Figure 2.**
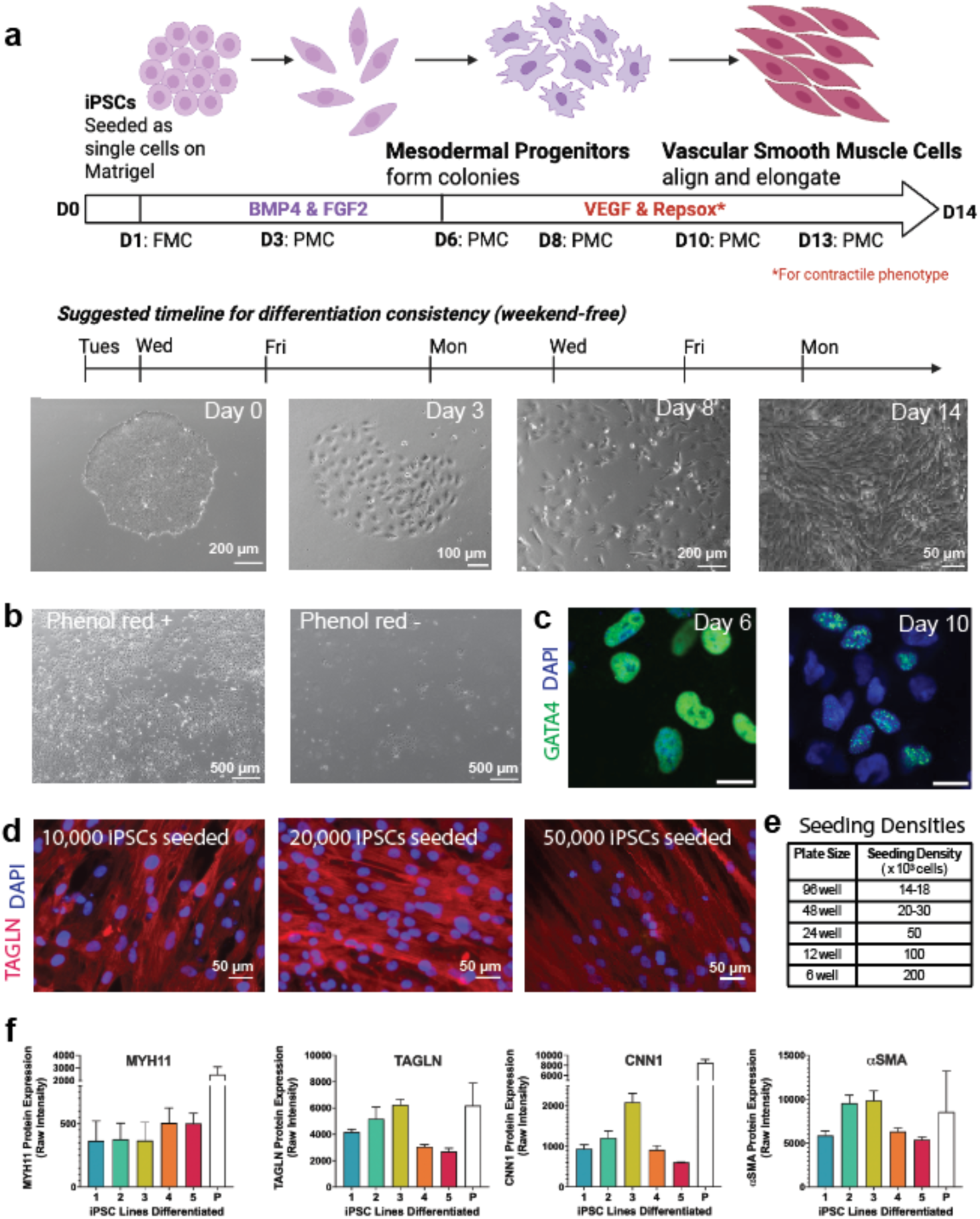
Representative mesodermal differentiation of iPSC lines. **a)** Schematic of the iVSMCs differentiation protocol describing full media change (FMC) and partial media changes (PMC), with suggested timelines to follow for weekend-free culturing, and with representative images of cells at day 0, 3, 8, 14; **b)** Phenol red-free DMEM/F12 media reduced the cellular yield of mesodermal progenitors; **c)** Expression of the early marker GATA4 in progenitor cells at day 6 was showed reduced expression of this early marker as cells begin to mature into VSMC-like cells on day 10. **d)** Cells seeded over a range of densities yielded more linear, elongated cells at higher density **e)** Cell seeding densitys were adjusted for plate size to account for the differences in tissue culture plates**. f)** Quantification of key VSMC contractile-state proteins expressed in iVSMCs (lines 1-5) compared to primary adult human aortic smooth muscle cells (P), as determined from proteomic analysis (raw intensity; n = 3-5 differentiations per line)

#### 2.2.1 WALL Media

Throughout the iPSC-derived modeling field, there is an increased demand for chemically-defined media due to the variable nature of animal-derived products, such as fetal bovine serum (FBS). The use of xenogenic-free reagents overcomes the possible development of xenogenic immune responses, and reduces animal to human transmittable diseases (xenozoonosis)^[26]^. Decreasing reliance on animal-based products in laboratory settings has also been encouraged following the COVID-19 pandemic, to further reduce risk of future zoonotic epidemics^[27]^. In vascular models, this practice is particularly important as FBS can alter VSMC extracellular matrix deposition and TGF-β signaling^[28]^. Luo and colleagues have previously demonstrated a xenogenic-free iVSMC differentiation protocol has no significant impact on the capacity of iPSCs to differentiate, moving the field closer towards clinical application of engineered VSMC cells^[29]^. Accordingly, here we have similarly employed a xenofree ‘WALL’ media comprising of DMEM/F12, insulin-transferrin-selenium, non-essential amino acids and GlutaMAX *in lieu* of undefined components (**Supplementary Table 1**). Interestingly, we observed a marked reduction in cell viability and proliferation towards mesenchymal progenitors using phenol red-free DMEM/F12, thus standard DMEM/F12 was utilized, although phenol red-free medium could be used in week 2 of the protocol (**Figure 2b, Supplementary Figure 1)**. This effect of phenol red-free DMEM/F12 may be associated with reduced estrogen signaling as the phenol-based dye is a known estrogen mimic which is known to influence proliferation in VSMCs^[30,31]^. Partial media changes were performed following an initial full media change to WALL media. While this is common practice in iPSC differentiation protocols in other fields^[32,33]^, it is not commonly stipulated in iVSMC protocols. Partial media changes capitalize on the growth factors and extracellular matrix (ECM) generated endogenously by the differentiating cells, and reduce effects of large fluctuations in osmolality and pH^[34]^.

To assess the effect of WALL media on protein expression of iVSMC compared to common commercially available media, proteomic analysis was performed on primary VSMC (human aorta-derived smooth muscle cells; HAoSMC) cultured in either WALL media or smooth muscle growth media 2 (SmGM-2, Lonza), for 8 days (**Supplementary Figure 2**). Cells cultured in either media exhibited similar morphology (**Supplementary Figure 2a**). WALL media-cultured cells proliferated were slower (**Supplementary Figure 2b**), and this more closely the turnover rates of quiescent contractile VSMCs *in vivo* that under normal physiological conditions turnover less than once per year^[35]^. Qualitative analysis of the proteomes of HAoSMCs cultured in WALL media compared to LONZA media indicated near identical protein expression, where 97.6% of the 3,443 proteins detected were in common between the two conditions (**Supplementary Figure 2c**). While While hierarchical analysis indicated sample clustering by media type (**Supplementary Figure 2d**), the overall proteomes of the WALL media-cultured cells were similar to those in commercial media, as indicated by similar Pearson’s correlation values between Lonza samples (0.870 – 0.926) and WALL samples (0.823 – 0.904), indicating consistent profiles between passages (**Supplementary Figure 2e**). Student’s t-test identified differentially expressed proteins between the two groups (**Supplementary Figure 2f)**, with differences primarily due to expression of proteins driven by the growth factors, IGF and PDGF, which are not utilized in the WALL medium differentiation (**Supplementary Figure 2g**).

#### 2.2.2 Initiation Format

To harness the ability of iVSMCs and their progenitors to synthesize biologically relevant ECMs *in situ*, and to ensure the protocol would lend itself to automation, we chose to use a monolayer iVSMC progenitor approach, rather than an embryoid body formation approach. Supplemention of the chemically defined media with BMP4 and FGF2 induced morphological changes indicative of mesodermal progenitor differentiation. Cells grew in colonies, prior to reaching confluence, and were GATA4 positive on day 6, with reduced expression by day 10 as they matured (**Figure 2c**). Cell seeding density was the largest contributor to cellular variability. Testing of different cell densities. (5 x 10^3^ cells/cm^2^, 12.5 x 10^3^ cells/cm^2^ or 25 x 10^3^ cells/cm^2^ in 24 well plates) indicated that cells grown at higher seeding densities had a more elongated and aligned morphology when stained for VSMC markers, including TAGLN, suggesting increased maturation in these cells (**Figure 2d**). Plate well-size also impacted differentiation, with smaller wells, such as those in 96-well plates, requiring a higher seeding density. This is likely because fluidic forces to which cells are exposed, including well surface area to volume ratios, base curvatures, and media handling differences, vary between tissue culture-plates of different sizes. Different seeding densities were thus established for different plate sizes, ranging from 21 - 56 x 10^3^ cells/cm^2^ (**Figure 2e**). Mid-differentiation passaging was avoided wherever possible, as repeated enzymatic digestion of the endogenously generated ECM markedly altered cell morphology and encouraged the development of the synthetic phenotype. If replating was required, this was performed on day 8 when the cells maintained more cellular plasticity. Confluence also varied markedly with substrate stiffness (glass compared to plastic), which has been previously reported, and is consistent with the ability of VSMCs to sense and respond to stiffness with shifts in proliferation^[10]^. To generate a robust culture of contractile iVSMCs, we employed Repsox, an inhibitor of TGF-β1 receptors^[36]^. Though many iVSMC protocols commonly use PDGF-BB and TGF-β1, we, similar to others^[37,38]^, observed that the use of these proteins increased the presence of a synthetic phenotype. Use of VEGF alone generated a more diverse culture with both contractile and synthetic cells, which may be of interest in disease modeling, while the combination of VEGF and Repsox produced a robust contractile culture. Proteomic analysis of these cells confirmed robust expression of hallmark VSMC proteins including MYH11, α-SMA, TAGLN and CNN1 (**Figure 2f**).

#### 2.2.3 Protein Expression Consistent with Robust Differentiation of iVSMCs

To examine batch-to-batch variation, all five iPSC lines were differentiated concurrently, repeating the differentiations three to five times to examine protein expression via immunocytochemistry and proteomics. Immunostaining indicated the differentiated iVSMCs were positive for hallmark VSMC markers, and quantification of positive cells (minimum 12,000 cells analyzed per line for each marker, n=5 lines) indicated robust expression of MYH11 (97 ± 0.82 SD % positive cells) **Figure 3a**), TAGLN (97.59 ± 2.2 SD % positive cells; **Figure 3b**) and CNN1(93 ± 1.3 SD % positive cells; **Figure 3c**), also validating the proteomic identification of these markers (in **Figure 2f**).

**Figure 3.**
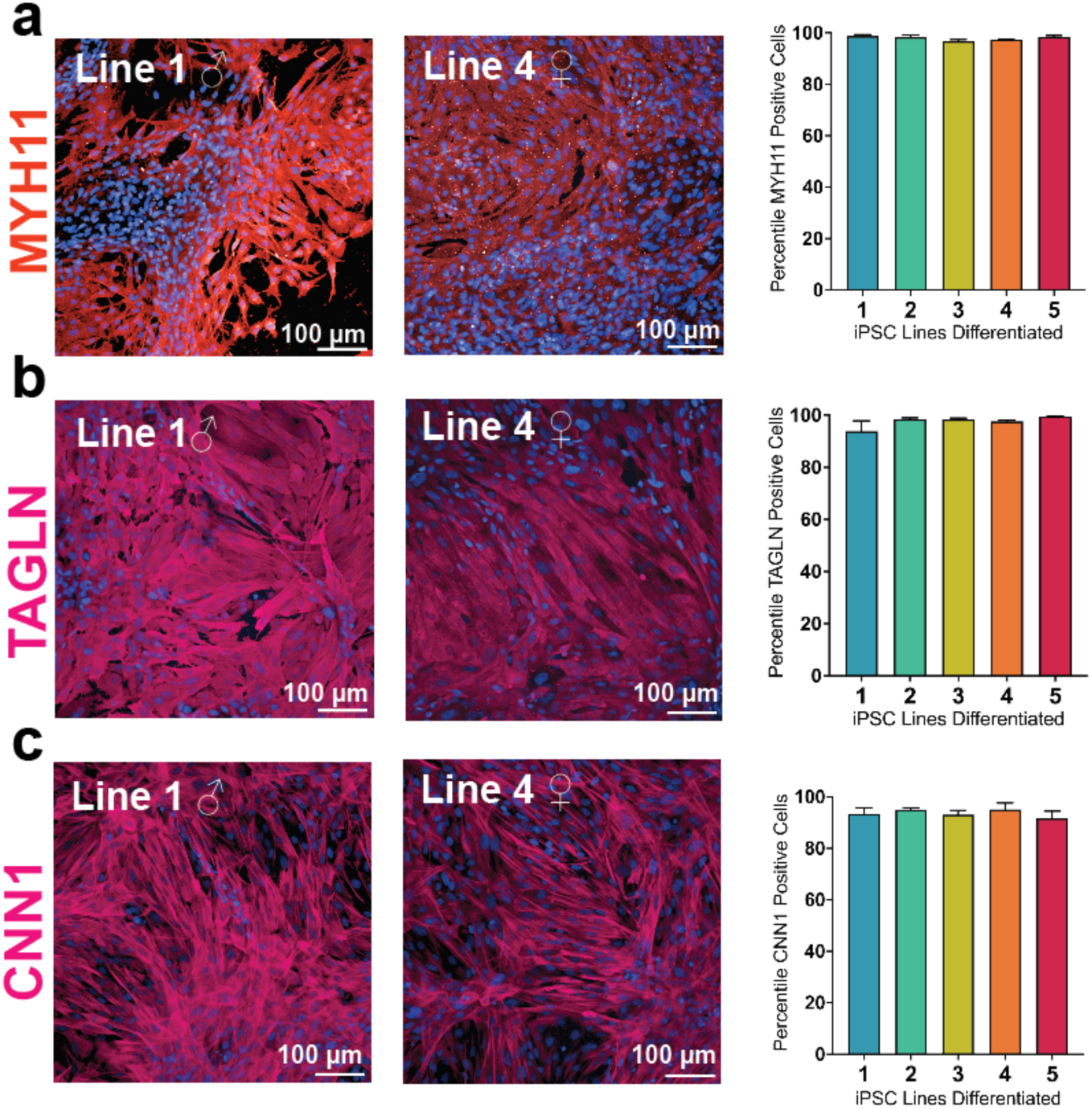
Representative Immunocytochemical Expression of VSMC Marker Proteins. **a)** MYH11, **b)** TAGLN and **c)** CNN1. Protein quantification was performed using ImageJ software applied to a minimum of 12,000 cells per line per marker in n= 3-5 differentiations per line.

### 2.3 WALL Differentiated iVSMCs are Contractile

To assess iVSMC function, intracellular calcium dynamics and contractile responses to carbachol (2 mM) were evaluated (**Figure 4**). Time-lapse microscopy using phase contrast imaging of the iVSMCs (**Figure 4a),** revealed significant carbachol-induced contraction of cells over a 10 minute period, as compared to baseline and to vehicle controls (**Figure 4b, Supplementary Video 1**). Carbachol-treated cells exhibited a 16 - 27% decrease in cell area, consistent with the normal range of VSMC contraction under physiological conditions. Characteristic calcium transient alterations were observed within the first minute of carbachol treatment (**Figure 4c**). An example of an individual iVSMC shifting its contractile response from single calcium peaks to rolling oscillations is shown in **Figure 4d**. Individual traces demonstrated a fast initial calcium spike, followed by a plateau phase lasting several seconds, often taking >20 seconds to return to baseline. Quantification of the sum of Cal-520AM dye fluorescence area revealed a ∼7-fold increase with carbachol treatment (**Figure 4e**). These findings indicate that iVSMCs generated with the WALL protocol were responsive to vasoactive signals, a key feature of smooth muscle physiology.

**Figure 4.**
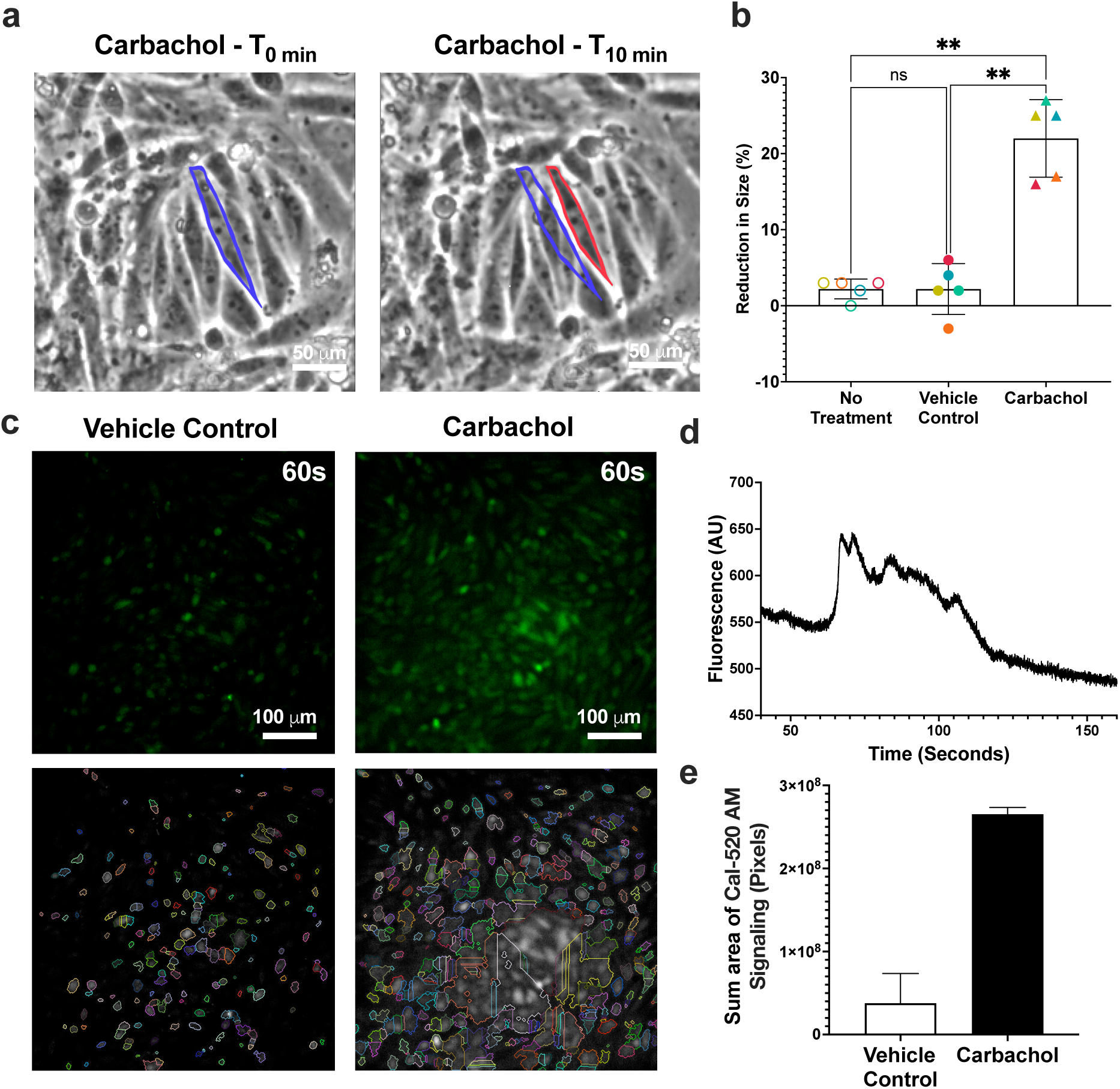
Functional characterization of iVSMCs. **a)** Representative phase contrast images of iVSMC pre- and post-carbachol treatment (2 mM, 10 min). **b)** Blinded single cell quantification of cell area prior to media change (no treatment), in response to vehicle control, and in response to carbachol treatment. Carbachol induced reductions in cells surface area by 16-27% (n= 5 lines, represented by colors, 10 cells/line; *:p < 0.01 determined via one-way ANOVA with Tukey’s post hoc test) indicating their contractile competency. **c)** Fluorescence imaging of Cal 520 AM dye-treated iVSMCs following vehicle (control) or carbachol treatment(2 mM, 1 min; top panels) and Region of Interest detection analysis via custom Python script of these areas (bottom panels). **d)** iVSMC calcium transients quantified via MATLAB. **e)** Sum of the area of fluorescence detected via Cal 520 AM imaging using a Nikon Eclipse Ti2-E Inverted Microscope.

### 2.4 iVSMC Proteome Unaltered by Robotic Media Changes

We next investigated the effects of using a large-scale liquid handling robotics system, Hamilton MicroLab STAR liquid handling robot (**Figure 5a**), for carrying out the differentiation procedure and performed comparisons with concurrently cultured, manually-differentiated cells. Gross morphologies were not altered by robotic liquid dispensing as observed by brightfield imaging (**Figure 5b**).

**Figure 5.**
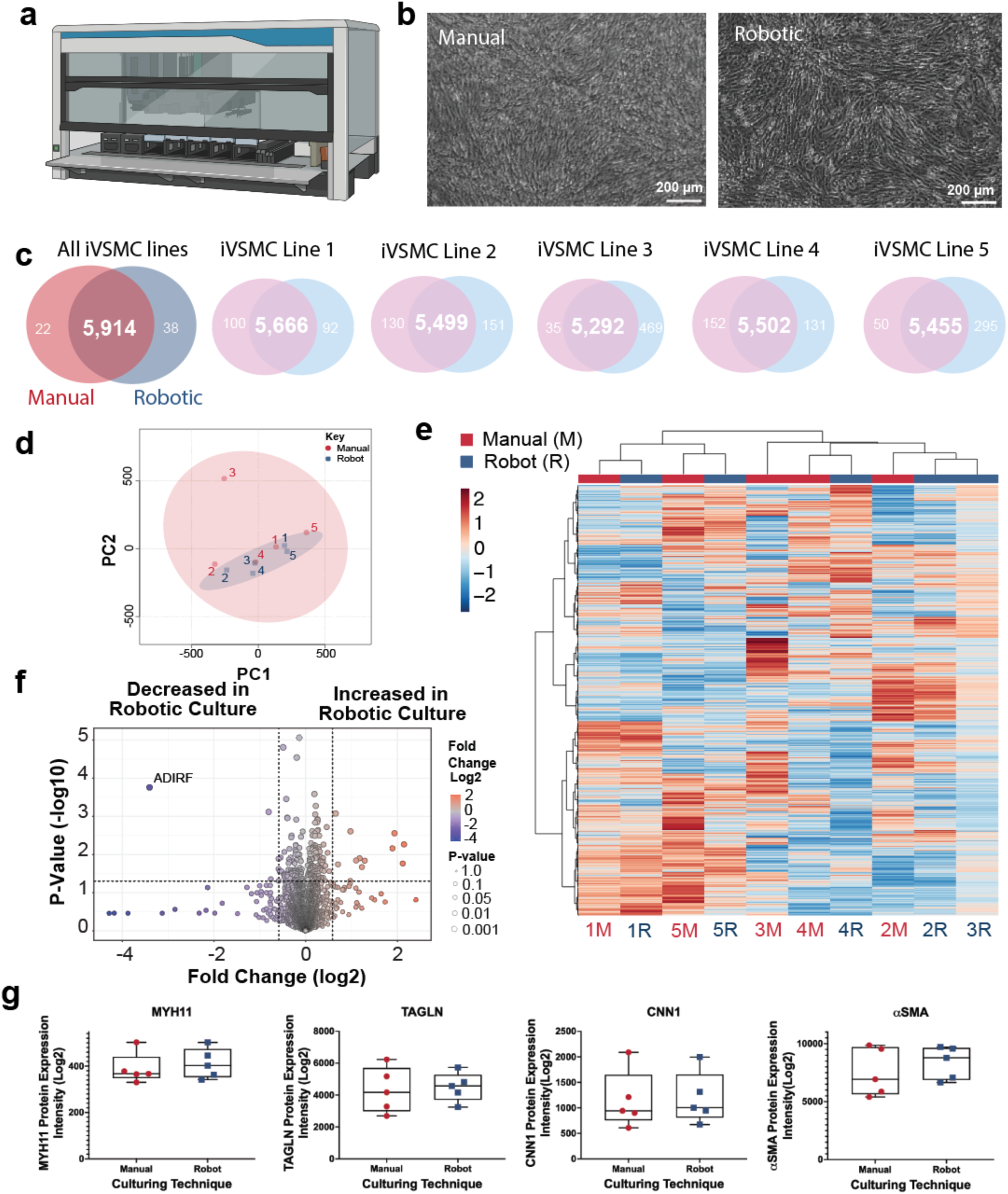
The iVSMC Proteome is Unaltered by Automated Robotic Liquid Compared to Manual Handling. **a)** iVSMCs differentiation was tested for feasibility of automation using a Hamilton MicroLab STAR Liquid handler coupled to an incubator system. **b)** Representative brightfield imaging using Incucyte shows similar morphology of cells cultured manually and robotically. **c)** Qualitative Venn diagram comparisons of the mean proteomes of five iVSMC lines (mean of n=5 lines differentiated concurrently, with n=5 consecutive differentiation replicates) either manually, or via robotic handling. **d)** With the exception of one outlier ( line 3 sample), principal component analysis clustered the average of samples by cell line (indicated by number) rather than by culture technique (manual (M) in red, robotic (R) in blue). **e)** Hierarchical analysis of the iVSMC proteomes indicated clustering of samples based on the cell lines (indicated by number) rather than the culture technique – Robotic (R) or manual static culture (M). **f)** Volcano plot representation of t-test scores indicates that few proteins are differentially expressed. **g)** Expression of the VSMC marker proteins, MYH11, TAGLN, CNN1 and aSMA was not different between cells cultured using robotic vs manual techniques.

**Figure 6.**
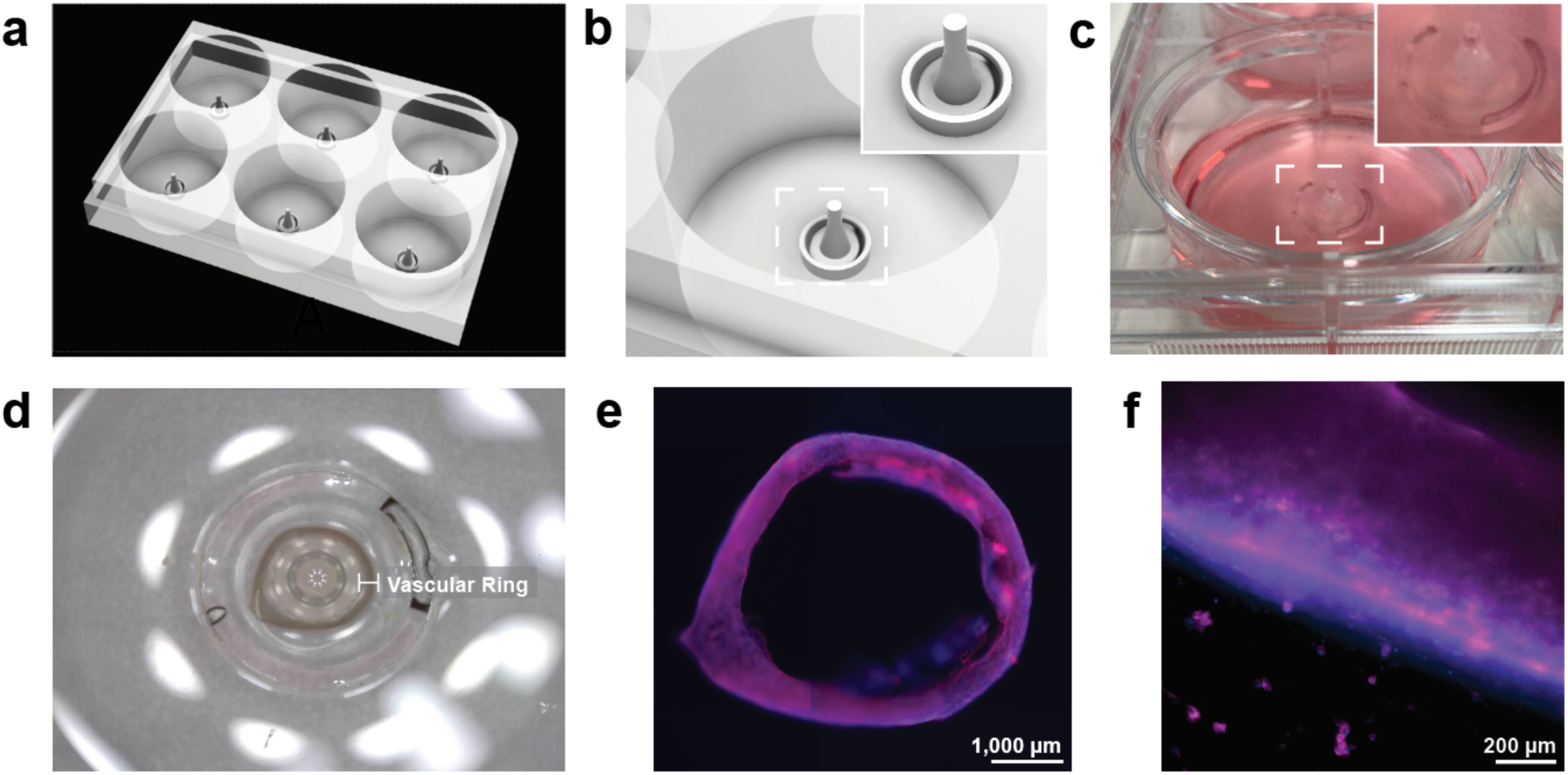
Differentiated iVSMCs were suitable for tissue engineering purposes. **a)** Bespoke vascular ring templates were designed via Fusion360 to generate vascular rings in a 6-well format to accommodate the same liquid handling workflow. **b)** The design with a tapered center to facilitate removal of ring for future applications. **c)** Polydimethylsiloxane molds were cast with polylactic acid. **d)** iVSMCs formed micro-tissues of sufficient integrity to be handled with tweezers for removal for downstream assays on day 4. **e)** Representative ring, approximately the size of a single well in a 96-well plate and **f)** overlay of immunocytochemical analysis of a ring showing positive expression of TAGLN (red), as well as DAPI (blue) staining, confirmed using a Celigo imaging cytometer connected to a Hamilton Robotics system.

To determine if robotic differentiation induced global changes, proteomic analyses was performed. iVSMCs from the five iPSC lines were differentiated manually or robotically in parallel across five sequential differentiations to ensure reproducibility and robustness. Proteins detection was highly consistent between the robotically cultured and manually cultured cells, with only 1.00% of proteins undetected between these culturing techniques (**Figure 5c).** The quantified proteomes consisted of 6,003 robustly detected proteins, including classic VSMC-associated proteins such as αSMA, which was the tenth most abundant protein (average of n=5 lines, 2 culture conditions), as well as MYH11, TAGLN, and CNN1, which were also confirmed by immunocytochemistry (Figure 3. Variability of marker concentration between differentiations was relatively low between cell lines (**Supplementary Figure 3**).

Principal component analysis of the cell line averages for each condition did not reveal partitioning based on culturing techniques, with cell line identity driving tight clustering in four of the five lines (**Figure 5d**). Hierarchical clustering similarly indicated that sample partitioning was determined by cell line, not culturing technique (**Figure 5e**). Comparison of the robotic and manual culturing of iVSMC Student’s t-test revealed no significant differences between robotic and manual iVSMC cultures using standard thresholds (fold change ≧2, FDR 0.1). Even with relaxed thresholds (fold change ≧1.5, p ≤0.05), only 18 proteins (0.29%) were differentially expressed between the two conditions (**Figure 5f; Supplementary Figure 4**). STRING pathway analysis revealed no pathway level enrichment between these 18 proteins (protein-protein interaction enrichment p-value = 0.89). No significant differences were detected in the hallmark VSMC proteins, MYH11, α-SMA, TAGLN and CNN1 (**Figure 5g).**

### 2.5 iVSMCs Capable of Tissue Engineering

To investigate the application of these cells in a tissue engineering setting, we utilized iVSMCs differentiated via automation to generate vascular rings. Using custom-designed polydimethylsiloxane (PDMS) ring-molds, we generated vascular ring tissues by combining Matrigel (∼10 mg/mL) and laminin (1 mg/mL) with iVSMCs (1 x 10^7^ cells/mL) dissociated on day 8 of the protocol, at a ratio of 1:1:1. Interestingly, resuspension in Matrigel alone was not sufficient to maintain the integrity of the vascular ring structures. Vascular rings were cultured for 4 days, at which time they held enough structural integrity to be removed from the pillar using tweezers. Immunocytochemical analysis indicated cells within the ring were positive for TAGLN.

#### Conclusion

In this study, we showcase a WALL protocol – a weekend-free, automatable, low-cost, low-variation protocol - to differentiate iPSCs into VSMCs. Using this protocol, we demonstrated for the first time that differentiation was unaffected by robotic liquid handling at the protein-pathway or functional level, addressing a critical bottleneck for the manufacturing of iVSMCs for tissue engineering and drug screening. Applied across a diverse cohort of iPSC lines from male and female donors of different ages, this method consistently yielded high-quality iVSMCs suitable for tissue engineering applications. The implementation of a chemically-defined differentiation medium marks a critical step toward a fully xenogenic-free protocol—an important advancement for improving biological relevance and in alignment with good manufacturing practice (GMP) standards required for translational and clinical applications. Importantly, we demonstrate for the first time that automation via robotic liquid handling does not introduce significant changes in protein expression or pathway-level interactions when compared to manual differentiation. These findings underscore the scalability and reproducibility of the WALL protocol, establishing a strong foundation for the standardized, large-scale generation of iVSMCs. This enables not only tissue engineering and regenerative medicine applications but also high-throughput drug screening and integration into vascular organ-on-a-chip models.

## 2. Experimental Section

### iPSC Reprogramming from Fibroblasts and Peripheral Blood Mononuclear Cells

All assays were conducted in compliance with the St. Vincent’s Hospital Human Research Ethics Committee (HREC/16/SVH/338, protocol number SVH 16/245). Cells were cultured at 37°C, under normoxic conditions (5% CO_2_). All reagents were purchased from ThermoFisher Scientific unless specified. Peripheral blood mononuclear cells (PBMCs) were reprogrammed as described previously^[19,21]^. Briefly, PBMCs were obtained from local pathology laboratories, following separation by Ficoll centrifugation, the cells were cultured in StemSpan erythroid progenitor medium (Stem Cell Technologies; SCT) according to the manufacturer’s guidelines. PBMCs at a density of 1×10^5^ were transduced using Cytotunes 2.0 Sendai Virus. After an incubation period of 24 hours, the cells were centrifuged, a full media change was performed, and cells were seeded in a single well of a 12-well tissue culture plate. After four days of culture, the cells were transferred to Matrigel (Corning) coated plates. The culture medium was supplemented with ReproTeSR (SCT), maintaining a 50:50 media ratio, and daily medium changes were conducted. Fibroblast samples were obtained from a skin punch biopsy, and reprogrammed as previously described^[39]^. The iPSC colonies that emerged were manually selected based on their characteristic iPSC morphology starting from day 15.

### Clonal Characterization

Clonal characterization was carried out as previously described^[40]^ unless otherwise stated. Briefly, iPSCs were first screened for morphology indicative of iPSCs, and Sendai Virus presence via PCR. Karyotyping was carried out by Monash Health Services, Victoria (Lines 4 and 5), or by Sullivan Nicolaides Pathology, Queensland. For all karyograms, a minimum of 15 cells were viewed at a resolution of 400 bands per haploid set by a cytogeneticist. Expression of pluripotency markers was determined immunocytochemically and by mRNA expression, using the Pluripotency Scorecard assay. Capacity for trilineage differentiation of the selected representative clones was similarly determined by protein and mRNA expression. iPSCs were maintained using ReLeSR passaging, seeding on Matrigel-coated plates in mTeSR Plus media.

### Differentiation to Synthetic and Contractile Phenotype VSMCs

iPSCs were single cell dissociated with Accutase (SCT) and plated at a density of 21 - 56 x 10^3^ cells/cm^2^ onto Matrigel with Y-27632 and supplemented with mTeSR Plus media overnight. A full media change using a chemically defined media (WALL media; Dulbecco’s modified eagle medium/nutrient mixture F-12, GlutaMAX, MEM non-essential amino acids, penicillin-streptomycin, insulin-transferrin-selenium) supplemented with bone morphogenetic protein 4 (BMP4) (50 ng/mL; SCT) and fibroblast growth factor 2 (FGF2) (20 ng/mL; SCT), was performed on day 1, and a partial media change was performed on day 3 for mesodermal progenitor generation. A partial media change was also performed on days 6, 8, and 10, using WALL media supplemented with vascular endothelial growth factor (VEGF) (50 ng/mL) and Repsox (25 µM).

#### Robotic Liquid Handling Differentiation

Simultaneous differentiation via robotic liquid handling and manual handling was conducted using the Hamilton MicroLab STAR liquid handling robot. All scripts and configurations used to run the Hamilton system are publicly available via Zenodo (DOI: 10.5281/zenodo.17355820).

### Immunocytochemistry and imaging

Adherent cells were processed as described^[32]^, including a 4% paraformaldehyde fixation (10 min, room temperature; RT), permeabilization (12 min, 0.1% Triton X /PBS) and 10% goat serum block for 1h at room temperature, and then incubated overnight with antibody (**Supplementary Table 2**) .Vascular rings were fixed for 20 min and then permeabilized for 20 min using 0.1% Triton X/PBS. Samples were washed with Tween 0.1% in PBS to remove residual primary antibody and incubated with a corresponding secondary antibody (1h RT), and DAPI stained. Fluorescence microscopy was performed as indicated using an EVOS cell imaging system (ThermoFisher), or a Nikon Eclipse Ti2-E inverted microscope, or Celigo (Revvity) imaging cytometry. Characterization of differentiation efficiency was quantified by unbiased imaging using an automated confocal microscopy (Opera Phenix, Revvity). Confluence and live cell tracking was performed using an in-incubator imager (Incucyte). Images were processed (overlay, scale bar addition, crop) with Image J.

### Mass Spectrometry Sample Preparation

Cells were harvested in RIPA buffer supplemented with protease inhibitor (cOmplete protease inhibitor; Roche) following aspiration of media and subsequent washes with PBS. Lysate was stored at -80°C. Cells were then lysed by thawing and repeated trituration through a 26.5 G needle, prior to centrifugation (20,000 x g for 30 min at 4°C) and harvesting of the supernatant fraction. Protein levels were determined using a BSA assay in accordance with the manufacturer’s guidelines (Pierce Rapid Gold BCA Protein Assay Kit #A53226, ThermoScientific, USA). Protein samples (100 µg in 100 µL of conditioned media) were prepared using the EasyPep Mini MS Sample Prep Kit (#A53226), following the instructions provided by the manufacturer. This process included a pre-digestion treatment with PNGaseF, a deglycosylating enzyme, before overnight trypsin digestion (ThermoFisher Scientific, USA). The resulting clean peptide samples were then dried by vacuum centrifuge and subsequently reconstituted in 100 µL of 0.1% formic acid in water, in preparation for LC-MS analysis.

### Label-free bottom-up quantification via micro-high-performance liquid chromatography coupled with quadrupole time-of-flight mass spectrometry (micro-HPLC-qTOF-MS)

LCMS data was acquired according to a previous study with some modifications in the liquid chromatography^[41]^. Briefly, a micro liquid chromatography system, Waters M-Class, coupled to SCIEX^TM^ 6600 Triple-TOF® mass spectrometer operating in positive electron spray ion mode (ESI +), was used to analyse the tryptic peptides within 33 minutes of acquisition. 4 µg of the peptide digest was injected onto nanoEase M/Z HSS T3 1.8μm 300μmx150mm (Waters, 186009249) column coupled to a Zorbax 300SB-C18, (5um, 5x0.3mm) guard column (Agilent, USA). The column temperature was maintained at 40 °C, and 98% water (2% acetonitrile) and acetonitrile containing 0.1% Formic acid were used as mobile phase A and B, respectively, at 5 µL/min flow rate with initial loading and later column cleaning and equilibration at 7µL/min. Briefly, the gradient started at a 7 µL/min flow rate at 2-10% B for 1.66 min and inclined to 10-25% B at 1.67-21.67 min at 5 µL/min flow rate, then 40-95% B at 23.33-24.67 min and kept for 2 minutes, then equilibrated for 9 min with 2% B at 7 µL/min flow rate. The TripleTOF® 6600 system was equipped with a DuoSpray source and 25 μm internal diameter electrode and controlled by Analyst 1.8.1 software and Analyst Device Driver for LC control. Ion source and MS parameters were set as follows: GS1 25; GS2 15; curtain gas 20; ion source temperature 150; ionSpray voltage floating of 5500; lower m/z limit 350; upper m/z limit 1250; window overlap (Da) 1.0; collision energy spread was set at 5. The Sequential windowed acquisition of all theoretical fragment ion spectra (SWATH™) acquisition method involved an MS1 scan ranging from m/z 350 to m/z 1,250 with an accumulation time of 50 ms, followed by 40 MS2 scans, each with a 35 ms accumulation time The MS2 scans utilized a variable precursor isolation width covering the m/z range of 400 to 1,250 (Supplementary Table 4). MS2 spectra were collected in the range of m/z 100 to 2000 in high-resolution mode, with ∼1.5 s total cycle time. PepCalMix calibrant (SCIEX, P/N 5045759, 10fmol/µL solution that was prepared by 1:100 dilution of the stock solution in dilution buffer consisting of 5% acetic acid and 2% acetonitrile) was injected every 12 samples for mass calibration along with the pooled quality control sample to evaluate the whole batch.

Six pooled QC samples were injected to generate DIA-only library using gas phase fractionation workflow^[42]^ with the following *m*/*z* mass ranges: 400–500, 500–600, 600–700, 700–800, 800–900, 900–1000, and 1000–1250. The precursor isolation window was set to *m*/*z* 5, with collision energy spread of 5 eV except for 700-990 and 990-1000 was 8 and 10 eV, respectively (**Supplementary Table 5**). The cycle time was 2.14 s, consisting of high- and low-energy scans with 40 ms MS2 accumulation time, and data were acquired in ‘high-resolution’ mode.

### Data processing and availability

Data were processed using Spectronaut v19.5 where DirectDIA+ analysis using BGS Factory workflow was implemented using canonical human FASTA database imported from Uniprot (25^th^ September 2024; 20,420 entries). Briefly, Pulsar search was done using the following parameters trypsin/P and LysC/P cleavage rules, specific digest type with 7-52 peptide length and max 2 missed cleavages. Carbamidomethyl (C) was set as a fixed modification and Acetyl (protein N-term), and oxidation (M) were set as variable modification with max variable modification set to 5 and identification FDR of 1% for PSM, peptide and protein group were applied for directDIA+ workflow with Automatic LFQ protein quantification method using the areas at MS2 level and automatic normalisation strategy. Automatic protein inference workflow using IDPicker algorithm was used for protein inference.

The raw and processed data have been deposited to the ProteomeXchange Consortium *via* the PRoteomics IDEntifications (PRIDE) repository^[43]^ with the dataset identifier PXD070184 and 10.6019/ PXD070184.

## Data analysis and Network Analysis

Relative protein intensity was statistically analyzed via metaboanalyst^[44]^. Data was log transformed, and Pareto scaled. Pearsons’s correlation of samples, hierarchical clustering heatmaps of normalised data (Standardization: Auto-scale features; Distance measure: Euclidean, Clustering method: Ward) and volcano plots were all generated with the platform. Fold change and multiple comparison adjusted P-value were as per in-text description. To analyse pathway-level interactions, Cytoscape software (version 3.8X) was implemented, including network clustering using Markov Cluster (MCL).

## Calcium Imaging

To measure calcium transients, iVSMCs were treated with Cal 520 AM dye (21130, AAT Bioquest) at a concentration of 5 μM for 30 min at 37°C. Dye was aspirated and cells were returned to iVSMC media (phenol red-free DMEM/F12) and allowed to equilibrate for 15 min. Cells were imaged in a live cell chamber (37°C and 5% CO_2_) using a Nikon Eclipse Ti2-E inverted microscope, utilizing a Nikon Plan Fluor 10× objective (NA, 0.3), and images were captured using an Andor Zyla sCMOS (Oxford Instruments). Cells were imaged for 10 minutes at baseline, followed by a vehicle control media change and carbachol treatment (2 mM ab141354, Abcam). Data acquisition was performed with a temporal resolution of 20 ms. Data analysis was conducted using a custom MATLAB script, as previously described^[45]^ to establish traces.

Sum of signaling area was analyzed using a custom Python pipeline generated using Google Colab, with assistance of ChatGPT 4o. Pipeline development incorporated scikit-image, OpenCV, NumPy, SciPy, pandas, and matplotlib. Grayscale video frames were extracted after skipping the initial 10% of the sequence to avoid transient surges. A baseline image was generated by averaging multiple sampled frames. Regions of interest (ROIs) were identified using Otsu thresholding, Gaussian filtering, and distance-transformed watershed segmentation (scikit-image). Noise was excluded by applying a minimum area threshold. ROI boundaries were visualized using the find contours function and overlaid on grayscale images with randomly assigned colors. Mean intensity values were extracted for each ROI across the time course (OpenCV, NumPy), normalized to baseline (F/F₀), and exported alongside the summed intensity signal for all ROIs per frame.

## Cell Size Quantification

iVSMC contractions were imaged using a Nikon Eclipse Ti2-E Inverted Microscope (10X), capturing images every 10s for 10 minutes via Andor Zyla sCMOS. Contractility was measured using Image J using a standard measure analysis at T0 and T10, randomly selecting cells. To measure cell size of HAoSMC, a standard measuring analysis using image J analysis was also performed. For these media comparisons, images of iVSMCs were taken using an EVOS brightfield microscope (ThermoFisher Scientific, USA) and blinded images were adjusted using Brightness and Contrast, and Threshold settings to outline cells. n=20 cells were catalogued as Regions of Interest and measured per image.

## Fabrication of iVSMC Vascular Rings

Master molds (negatives) for fabricating PDMS pillars to support ring formation were 3D printed using an Asiga Max X35 (365 nm) stereolithography printer using Asiga DentaMODEL resin. Print parameters were set to 25 μm layer thickness, 0.27 s exposure time, 32°C build temperature, and 30% light intensity. After printing, molds were first washed gently in 99% isopropyl alcohol for at least 1 hour on an orbital shaker to remove uncured resin, then UV cured for 3 × 15 minutes, and finally baked overnight at 65°C. PDMS was prepared by mixing Sylgard 184 (10:1 base:curing) and degassing in a vacuum chamber for 1 hour. Once ready, each well inside a 6 well plate (Corning) was cleaned with 100% ethanol and 2 mL of PDMS was pipetted into each well to form a thin layer of PDMS. This layer was then cured for 45 min at 65°C. This layer helps to ensure that molds detach without ripping off the surface of the well plate. Next, 3D printed molds were placed in an oxygen plasma chamber and exposed to plasma for 30 seconds, followed by pipetting of PDMS into each mold and degassing for 2 minutes to remove trapped air bubbles. The molds were then inverted and placed on top of the cured PDMS layers within each well of the 6-well plate. The assembled system was cured overnight at 65°C. After cooling, the 3D printed molds were carefully removed using pliers leaving behind the PDMS positives.

Tissues were generated by seeding 1 x 10^7^ iVSMCs on day 10 of the differentiation in 100 uL of Matrigel:laminin (1:1 ratio) into the ring wells and left to gel for 30 min prior to gently filling the well with media. Media changes were carried out as normal.

## Statistical analysis

Numerical data are presented as mean ± S.E.M., unless otherwise stated. Sample sizes and number of independent experiments are specified in figure legends. Graphical representations and statistical analyses were conducted using GraphPad Prism version 10. To assess the significance of differences between a control and experimental group, an unpaired two-tailed Student’s t-test was employed. For comparisons among multiple groups, one-way ANOVA was utilized with *post hoc* testing as described for each.

## Acknowledgements

The authors wish to acknowledge the generous contributions of our cell donors, the ongoing support of the SCADaddle Inc. group, and the valuable input of the VCCRI Arteriopathy-SCAD Cohort (VASC) team. We wish to thank Prof Joesph Wu, who kindly provided one of the iPSC lines used in this work. We also wish to thank Dr Osvaldo Contreras and Prof Richard Harvey for use of their tissue culture facilities. This work was partially supported by funding from the Australian National Health & Medical Research Council (APP1161200), NSW Health (Senior Clinician Cardiovascular Grant RG193092, and Senior Scientist Cardiovascular Grant RG04492) and a L3 National Health and Medical Research Council, Australia Investigator Grant (ID: 2020203) awarded to RMG; Animal component-free work was supported by a Medical Advances Without Animals Fellowship and an AMP Tomorrow Maker grant awarded to MB. Portions of this research were conducted using the Victor Chang Cardiac Research Institute Innovation Centre, established with funding from the New South Wales Government. JT and VR were supported by Office of Health and Medical Research Non-animal Technologies Network; NSW Cardiovascular Capacity building scheme collaborative research grant that supported JR, and MRFF Stem cell mission (2024443) also supported JT.

## Conflicts of Interest

The authors declare no conflicts exist.

---

**Supplementary Figure 1.**
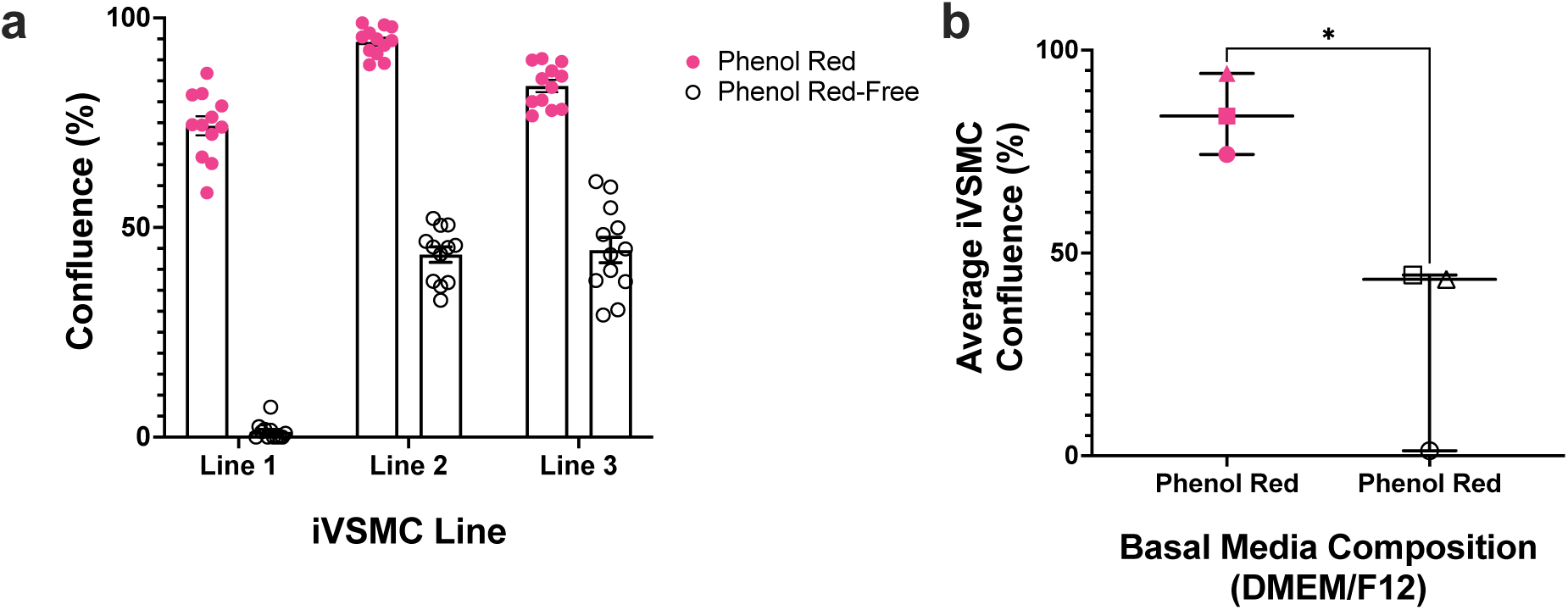
**Phenol red in iVSMC basal media influences differentiation to mesenchymal progenitors**. **A.** Differentiation of iPSCs (n=3 female-derived lines) towards mesenchymal progenitors was simultaneously performed in phenol red and phenol red-free DMEM/F12. Confluence was quantified on Day 3 of differentiation, via Incucyte live cell imaging, and analyzed via Incucyte Zoom software (n=12 technical replicates). **B.** Mean average confluences of these iVSMCs (lines represented by different symbols) indicated a significantly greater (Student’s t-test, *P = 0.032; n =3 biological replicates) confluence on Day 3 of iVSMCs differentiated in phenol red-containing medium compared to phenol red-free medium.

**Supplementary Figure 2.**
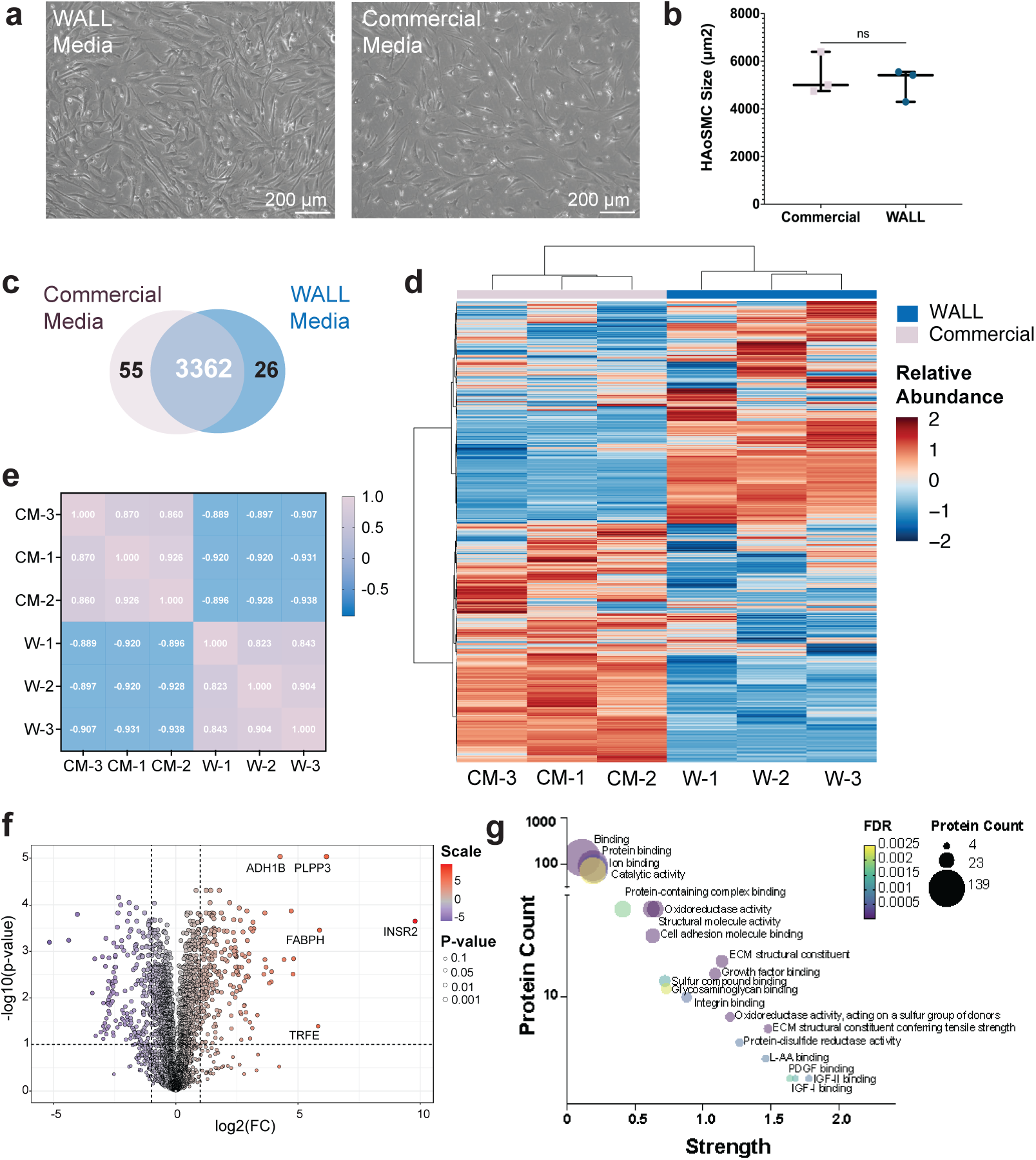
Comparison of primary iVSMCs Grown in WALL media (W) Commerical Media (CM) Comparison. **A.** Brightfield imaging indicates similar VSMC morphology in primary cells cultured in iVSMC media or commercially available media. **B.** Cell size analysis of HAoSMCs cultured in the two media types, as measured on day 5 post passage via Incucyte imaging and ImageJ analysis from consecutive weeks of culturing (n=3; blinded) **C.** Qualitative Venn diagram comparison of the mean proteomes of HAoSMCs cultured concurrently with Wall media or the common commercial media, LONZA. **D.** Hierarchical analysis indicates sample clustering based on media type, **E.** Pearson’s analysis. **F.** Volcano plot representation of t-test scores highlights differentially expressed proteins. **G.** Pathway analysis of the significantly different proteins highlights only minor differences that are growth factor-related.

**Supplementary Figure 3.**
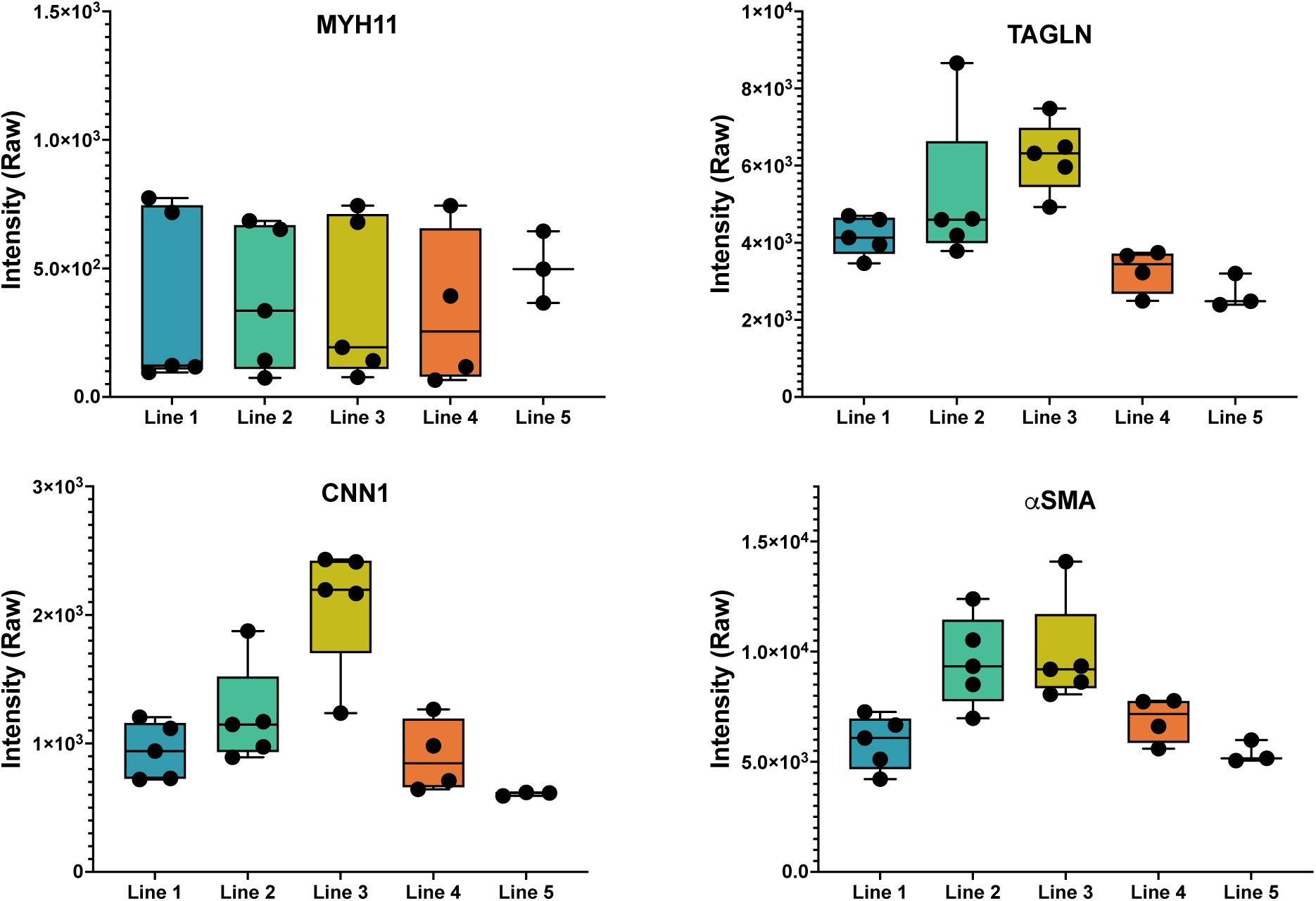
Consistency of hallmark VSMC markers MYH11, TAGLN, CNN1 and αSMA between differentiations of iVSMC cell lines. Values are raw intensity from proteomic quantification. Each datapoint represents a separate differentiation.

**Supplementary Figure 4.**
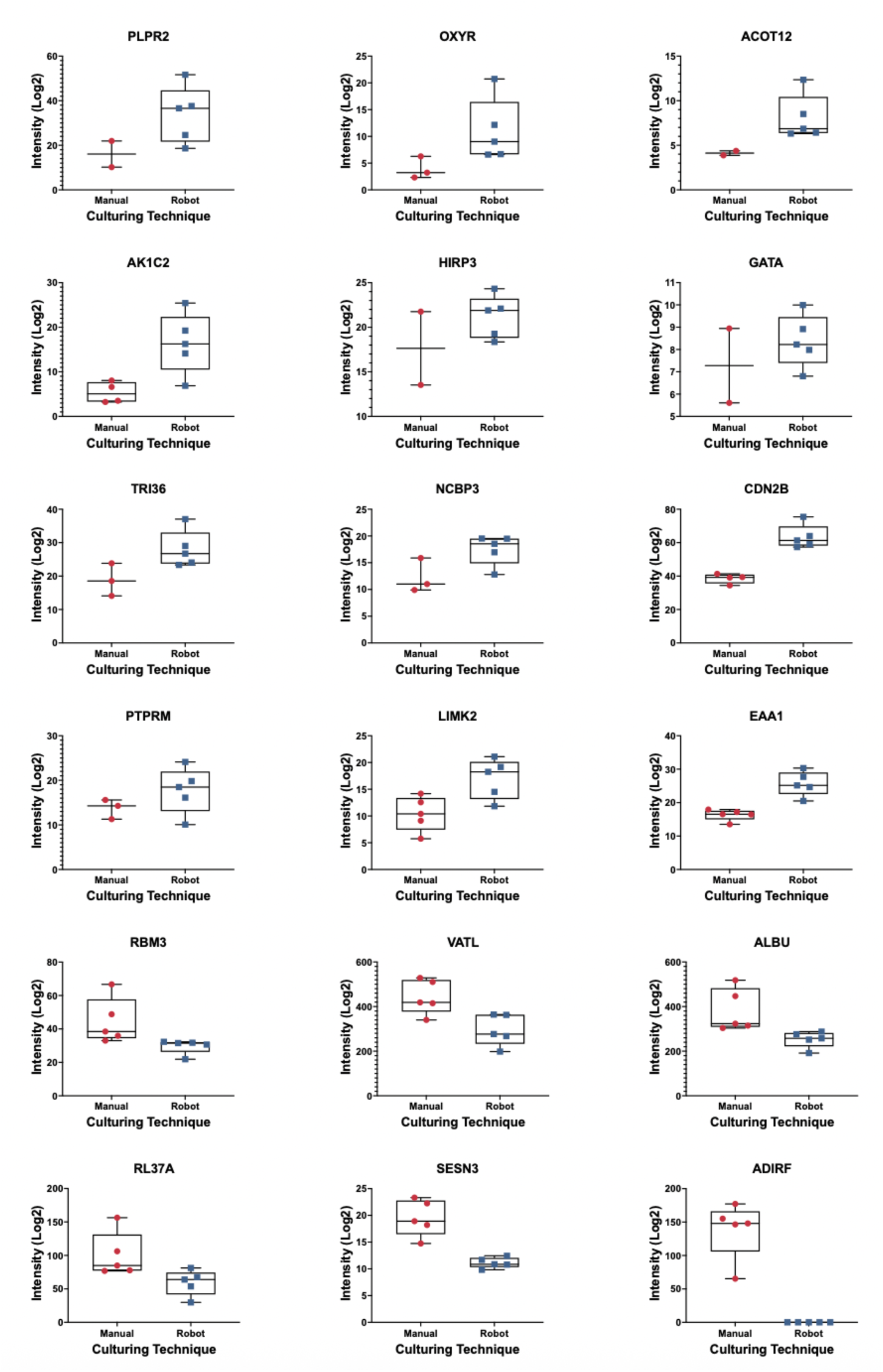
Proteins differentially expressed between manual and robot culturing of iVSMCs using relaxed thresholds. 18 proteins were detected as being differentially expressed with relaxed thresholding (1.5 fc, 0.05 Raw p-value).

**Supplementary Table 1.** WALL Media Composition

| **Component Shorthand** | **Component** | **Volume** | **Supplier** | **Catalogue Number** |
| --- | --- | --- | --- | --- |
| ITS | Insulin-transferrin-selenium | 5 mL | Life Technologies | 41400045 |
| MEM NEAA | Minimum Essential Medium Non-Essential Amino Acids | 5 mL | Life Technologies | 11140050 |
| GlutaMAX | GlutaMAX Supplement | 5 mL | Life Technologies | 35050061 |
| Optional: Penstrep | Penicillin-Streptomycin | 5 mL | Life Technologies | 15070063 |
| DMEM/F12 | Dulbecco's Modified Eagle Medium/Nutrient Mixture F-12 | 500 mL Total Volume | Life Technologies | 11320-033 |

**Supplementary Table 2.** – iVSMC and HAoSMC: CSV

**Supplementary Table 3.** – Proteomics: Static, normal culture compared to robotic culture

**Supplementary Table 4.**
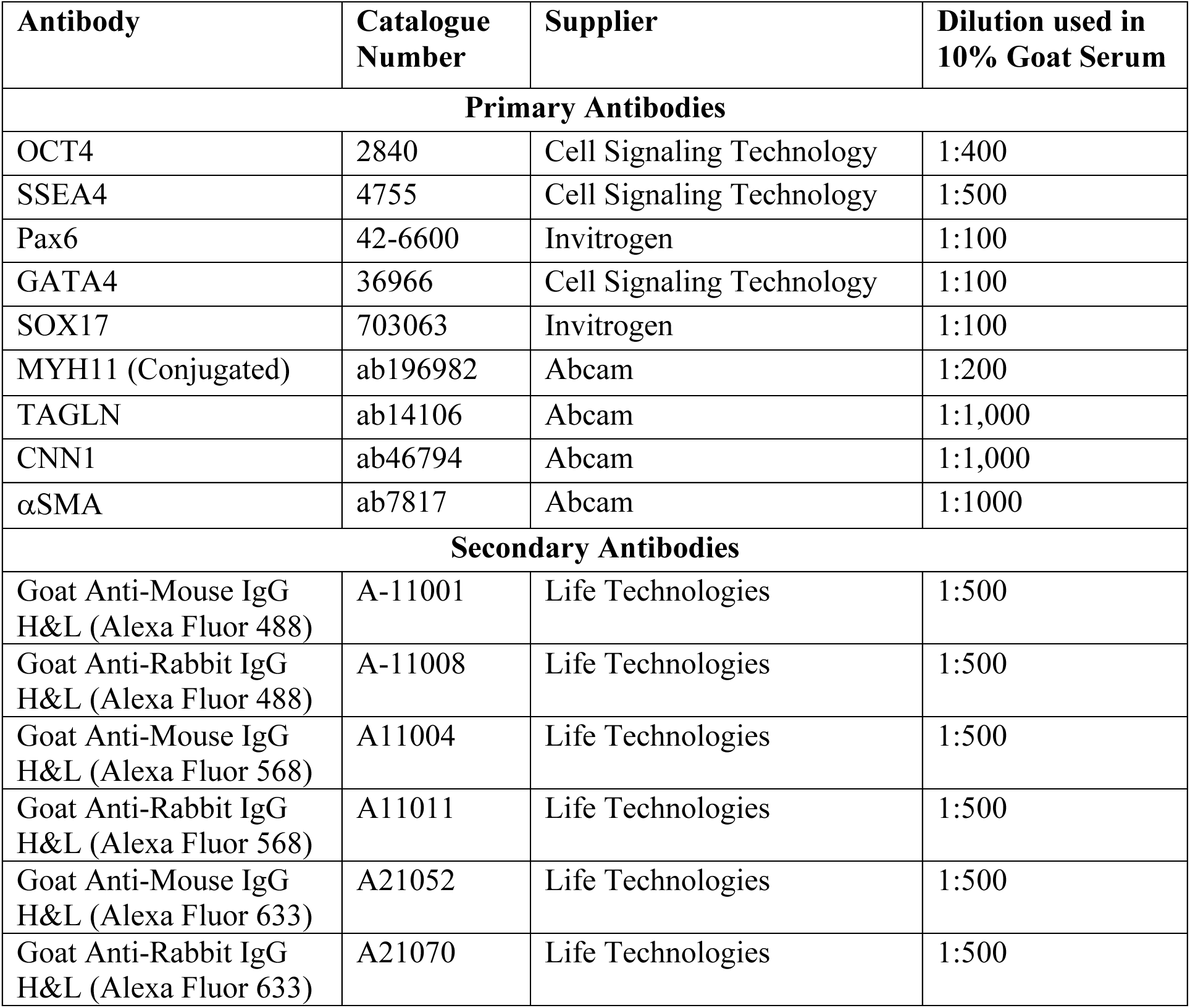
– Antibody Table

## Notes

### Competing Interest Statement

The authors have declared no competing interest.

## References

[1] American Heart Association, American Stroke Association, Cardiovascular Disease: A Costly Burden for America - Projections Through 2035, 2017.

[2] H. Wolinsky, S. Glagov, Circulation Research 1967, 20, 99.

[3] M. Bax, J. Thorpe, V. Romanov, Front. Sens. 2023, 4, 1294721.

[4] S. E. Iismaa, X. Kaidonis, A. M. Nicks, N. Bogush, K. Kikuchi, N. Naqvi, R. P. Harvey, A. Husain, R. M. Graham, Regen Med 2018, 3, 6.

[5] M. Bax, V. Romanov, K. Junday, E. Giannoulatou, B. Martinac, J. C. Kovacic, R. Liu, S. E. Iismaa, R. M. Graham, Front. Cardiovasc. Med. 2022, 9, 1055862.

[6] S. Allahverdian, C. Chaabane, K. Boukais, G. A. Francis, M.-L. Bochaton-Piallat, Cardiovascular Research 2018, 114, 540.

[7] G. L. Basatemur, H. F. Jørgensen, M. C. H. Clarke, M. R. Bennett, Z. Mallat, Nat Rev Cardiol 2019, 16, 727.

[8] S. Ayoubi, S. P. Sheikh, T. V. Eskildsen, Cardiovascular Research 2017, 113, 1282.

[9] V. Sorokin, K. Vickneson, T. Kofidis, C. C. Woo, X. Y. Lin, R. Foo, C. M. Shanahan, Front. Immunol. 2020, 11, 599415.

[10] P. Swiatlowska, B. Sit, Z. Feng, E. Marhuenda, I. Xanthis, S. Zingaro, M. Ward, X. Zhou, Q. Xiao, C. Shanahan, et al., Sci. Adv. 2022, 8, eabm3471.

[11] V. Hosseini, A. Mallone, N. Mirkhani, J. Noir, M. Salek, F. S. Pasqualini, S. Schuerle, A. Khademhosseini, S. P. Hoerstrup, V. Vogel, Adv. Sci. 2020, 7, 2070063.

[12] M. Shen, T. Quertermous, M. P. Fischbein, J. C. Wu, Circ Res 2021, 128, 670.

[13] C. S. Kwartler, J. E. E. Pinelo, n.d.

[14] S. Sinha, D. Iyer, A. Granata, Cell. Mol. Life Sci. 2014, 71, 2271.

[15] M.-N. Doulgkeroglou, A. Di Nubila, B. Niessing, N. König, R. H. Schmitt, J. Damen, S. J. Szilvassy, W. Chang, L. Csontos, S. Louis, et al., *Front. Bioeng. Biotechnol.* 2020, 8, 811.

[16] V. Hosseini, A. Mallone, N. Mirkhani, J. Noir, M. Salek, F. S. Pasqualini, S. Schuerle, A. Khademhosseini, S. P. Hoerstrup, V. Vogel, Advanced Science 2020, 7, 2000173.

[17] Z.-D. Shi, J. M. Tarbell, Ann Biomed Eng 2011, 39, 1608.

[18] N. Sun, M. Yazawa, J. Liu, L. Han, V. Sanchez-Freire, O. J. Abilez, E. G. Navarrete, S. Hu, L. Wang, A. Lee, et al., Sci. Transl. Med. 2012, 4, DOI 10.1126/scitranslmed.3003552.

[19] M. Bax, K. Junday, E. Hurley, S. Hesselson, L. McGrath-Cadell, X. Kaidonis, I. Tarr, E. Giannoulatou, S. E. Iismaa, R. M. Graham, Stem Cell Research 2025, 87, 103770.

[20] K. Mishra, K. Junday, C. M. Y. Wong, A. Y. Chan, S. Hesselson, D. W. Muller, S. E. Iismaa, A. Mehta, R. M. Graham, Stem Cell Research 2019, 41, 101584.

[21] M. Bax, K. Junday, S. E. Iismaa, X. Kaidonis, D. Muller, S. Hesselson, R. M. Graham, Stem Cell Research 2023, 73, 103238.

[22] R. J. G. Hartman, D. M. C. Kapteijn, S. Haitjema, M. N. Bekker, M. Mokry, G. Pasterkamp, M. Civelek, H. M. den Ruijter, Sci Rep 2020, 10, 12367.

[23] N. R., D. Gupta, S. Escopete, D. Rai, A. Stotland, N. Sundararaman, B. Ngu, K. Dabke, L. McCarthy, R. S. Santos, et al., IJMS 2024, 26, 187.

[24] V. Lo Sardo, W. Ferguson, G. A. Erikson, E. J. Topol, K. K. Baldwin, A. Torkamani, Nat Biotechnol 2017, 35, 69.

[25] Australian Bureau of Statistics, n.d.

[26] L. E. Chapman, T. M. Folks, D. R. Salomon, A. P. Patterson, T. E. Eggerman, P. D. Noguchi, N Engl J Med 1995, 333, 1498.

[27] S. O. Petrovan, D. C. Aldridge, H. Bartlett, A. J. Bladon, H. Booth, S. Broad, D. M. Broom, N. D. Burgess, S. Cleaveland, A. A. Cunningham, et al., Biological Reviews 2021, 96, 2694.

[28] B. Bilgen, E. Orsini, R. K. Aaron, D. McK. Ciombor, J Tissue Eng Regen Med 2007, 1, 436.

[29] J. Luo, Y. Lin, X. Shi, G. Li, M. H. Kural, C. W. Anderson, M. W. Ellis, M. Riaz, G. Tellides, L. E. Niklason, et al., Acta Biomaterialia 2021, 119, 155.

[30] W. V. Welshons, M. F. Wolf, C. S. Murphy, V. C. Jordan, Molecular and Cellular Endocrinology 1988, 57, 169.

[31] J. Song, Y. Wan, B. E. Rolfe, J. H. Campbell, G. R. Campbell, Atherosclerosis 1998, 140, 97.

[32] M. Bax, J. McKenna, D. Do-Ha, C. H. Stevens, S. Higginbottom, R. Balez, M. e C. Cabral-da-Silva, N. E. Farrawell, M. Engel, P. Poronnik, et al., Cells 2019, 8, 581.

[33] S. S. Muñoz, M. Engel, R. Balez, D. Do-Ha, M. C. Cabral-da-Silva, D. Hernández, T. Berg, J. A. Fifita, N. Grima, S. Yang, et al., Cells 2020, 9, 2018.

[34] C. Maass, M. Dallas, M. E. LaBarge, M. Shockley, J. Valdez, E. Geishecker, C. L. Stokes, L. G. Griffith, M. Cirit, Sci Rep 2018, 8, DOI 10.1038/s41598-018-25971-y.

[35] R. A. Neese, L. M. Misell, S. Turner, A. Chu, J. Kim, D. Cesar, R. Hoh, F. Antelo, A. Strawford, J. M. McCune, et al., Proc Natl Acad Sci U S A 2003, 99, 15345.

[36] J. K. Ichida, J. Blanchard, K. Lam, E. Y. Son, J. E. Chung, D. Egli, K. M. Loh, A. C. Carter, F. P. Di Giorgio, K. Koszka, et al., Cell Stem Cell 2009, 5, 491.

[37] J. Zhang, B. E. McIntosh, B. Wang, M. E. Brown, M. D. Probasco, S. Webster, B. Duffin, Y. Zhou, L.-W. Guo, W. J. Burlingham, et al., Stem Cell Reports 2019, 12, 1269.

[38] L. Liu, C. Jouve, J. Henry, T.-E. Berrandou, J.-S. Hulot, A. Georges, N. Bouatia-Naji, Hypertension 2023, 80, 740.

[39] M. Bax, R. Balez, S. S. Muñoz, D. Do-Ha, C. H. Stevens, T. Berg, M. C. Cabral-da-Silva, M. Engel, G. Nicholson, S. Yang, et al., Stem Cell Research 2019, 40, 101530.

[40] M. Bax, K. Junday, S. E. Iismaa, X. Kaidonis, D. Muller, S. Hesselson, R. M. Graham, Stem Cell Research 2023, 73, 103238.

[41] J. Muenzner, P. Trébulle, F. Agostini, H. Zauber, C. B. Messner, M. Steger, C. Kilian, K. Lau, N. Barthel, A. Lehmann, et al., Nature 2024, 630, 149.

[42] L. K. Pino, S. C. Just, M. J. MacCoss, B. C. Searle, Molecular & Cellular Proteomics 2020, 19, 1088.

[43] Y. Perez-Riverol, A. Csordas, J. Bai, M. Bernal-Llinares, S. Hewapathirana, D. J. Kundu, A. Inuganti, J. Griss, G. Mayer, M. Eisenacher, et al., Nucleic Acids Research 2019, 47, D442.

[44] Z. Pang, L. Xu, C. Viau, Y. Lu, R. Salavati, N. Basu, J. Xia, Nat Commun 2024, 15, 3675.

[45] J. Thorpe, M. D. Perry, O. Contreras, E. Hurley, G. Parker, R. P. Harvey, A. P. Hill, J. I. Vandenberg, Stem Cell Res Ther 2023, 14, 183.

